# Contextual and temporal regulation of fear memory consolidation in the basolateral amygdala complex

**DOI:** 10.1101/2022.12.03.518947

**Authors:** Jessica Leake, Luisa Saavedra Cardona, R. Frederick Westbrook, Nathan M. Holmes

**Affiliations:** School of Psychology, University of New South Wales, Sydney, NSW, Australia, 2052

**Keywords:** Context, fear conditioning, basolateral amygdala, protein synthesis, consolidation

## Abstract

It is widely accepted that fear memories are consolidated through protein synthesis-dependent changes in the basolateral amygdala complex (BLA). However, recent studies show that protein synthesis is *not* required to consolidate the memory of a new dangerous experience when it is similar to a prior experience. Here, we examined whether the protein synthesis requirement for consolidation of the new experience varies with its spatial and temporal distance from the prior experience. In each experiment, rats were conditioned to fear a stimulus (S1, e.g., light) across its pairings with shock in stage 1; and a second stimulus (S2, e.g., tone) that preceded additional S1-shock pairings (S2-S1-shock) in stage 2. The latter stage was followed by a BLA infusion of a protein synthesis inhibitor, cycloheximide or vehicle. Finally, rats were tested for fear to S2. Critically, protein synthesis in the BLA was *not* required to consolidate fear to S2 when the two training stages occurred 48 hours apart and in the same context; *was* required when the two training stages were separated by a 14-day delay or occurred in different contexts; but was *again not* required when rats were re-exposed to S1 or shock after the delay or in the different context. Thus, protein synthesis in the BLA is not always required to consolidate a new fear memory. Instead, this requirement is determined by the degree of similarity between present and past experiences, the time and place in which those experiences occur, as well as reminders of the past experience.

**Significance Statement:** Protein synthesis in the basolateral amygdala complex (BLA) is *not* required to consolidate the memory of a new dangerous experience when it is similar to a prior experience. This study is significant in showing that: 1) when the new, similar experience occurs after a delay or in a different context, the protein synthesis requirement for its consolidation is reinstated; and 2) the effects of the delay and context shift are reversed by reminding animals of their prior experience. Thus, the neural mechanisms underlying memory consolidation are dynamically regulated by similarity/dissimilarity between present and past experiences, the time and place in which those experiences occur, as well as reminders of the past experience.

## Introduction

People readily learn about stimuli that signal danger. In the laboratory, this type of learning is studied using Pavlovian threat or fear conditioning in rodents. In a standard protocol, rats are placed in a distinctive chamber where they receive pairings of an initially neutral conditioned stimulus (CS), such as a tone or a light, and an innately aversive unconditioned stimulus (US), such as foot-shock. Rats display defensive responses such as freezing when subsequently presented with the CS alone. These responses are taken to mean that the pairings produce an enduring memory of the association between the CS and US, which is retrieved by the CS and expressed in defensive responses indicative of fear in people.

The consolidation of a CS-US association in long term memory depends on cellular processes in a network of brain regions, including synthesis of new proteins in the basolateral amygdala complex (BLA). This is supported by findings that direct infusion of a protein synthesis inhibitor into the BLA immediately before or after a conditioning session spares the acquisition of fear responses to the CS but impairs their expression when the CS is tested days later (Schafe and LeDoux, 2000; Maren et al., 2003; Desgranges et al., 2008). Such findings have been taken to mean that new proteins are essential for stabilising learning-induced changes in BLA neurons. However, these findings have emerged from studies involving a single fearful experience (i.e., acquisition of fear to a CS paired with a US), and recent evidence shows that the molecular mechanisms that support a second fearful experience may differ from those that support an initial fearful experience as a function of their similarity.

This evidence comes from studies that have used a two-stage training protocol (Leidl et al., 2018; Williams-Spooner et al., 2019). In stage one, rats were exposed to pairings of a novel stimulus (S1; e.g., tone) and shock, which resulted in formation of an initial S1-shock fear memory. In stage two, rats were exposed to a second novel stimulus (S2; e.g., light) in sequence with S1 and shock (S2-S1-shock sequence), which resulted in formation of a second fear memory involving the S2. Intra-BLA infusions of the protein synthesis inhibitor, cycloheximide before or after stage two had no effect on the new S2 fear memory. By contrast, a BLA cycloheximide infusion disrupted fear to S2 among rats exposed to S2-S1-shock sequences in the absence of any prior training; and among rats exposed to S2-[trace]-shock pairings in stage two after having been exposed to S1-shock pairings in stage one. Together, these findings show that the protein synthesis requirement for consolidating fear to S2 can be influenced by the prior conditioning of S1: specifically, when S1 is conditioned in stage 1 and present in stage 2, consolidation of fear to S2 does not require protein synthesis in the BLA.

The present series of experiments used the two-stage conditioning protocol described above to examine whether context and time regulate the protein synthesis requirement for consolidating fear to the S2. It focused on context and time as contextual- and temporal-dependence are hallmarks of episodic memory (Tulving, 1972) and both factors critically influence memory encoding and recall (Goodwin et al., 1969; Godden and Baddeley, 1975; Kahana, 1996; Yonelinas et al., 2019). However, it is unknown whether they influence the molecular mechanisms of memory consolidation in the BLA. The initial experiments replicated the previous finding that, when the session of S2-S1-shock sequences occurs 48 hours after the session of S1-shock pairings and in the same context, consolidation of fear to the S2 does not require protein synthesis in the BLA. Subsequent experiments then examined whether this protein synthesis requirement is reinstated when the two training stages are separated by 14 days or administered in different physical contexts; and whether any reinstatement can be reversed or blocked by reminding rats of their stage one training after the 14-day delay or in the different context.

## Materials and Methods

### Subjects

Subjects were 239 experimentally naïve adult, male and female Long Evans rats, obtained from the breeding facility at the University of New South Wales. Male and female rats were housed separately in plastic tubs (22 cm high x 67 cm long x 40 cm wide) with four males and up to eight females per tub. Rats were housed in a temperature-controlled colony room (22°C) on a 12-hour light-dark cycle (lights on at 0700) with continuous access to food and water. Rats were handled once a day for at least three days prior to beginning behavioral procedures.

### Apparatus

Behavioral procedures took place in two distinct contexts referred to as contexts A and B. Context A consisted of four identical chambers (30 cm high x 26 cm long x 30 cm wide) with a grid floor composed of rods that were seven mm in diameter, spaced 18 mm apart. Context B was located in a different room and consisted of four identical chambers (30 cm high x 26 cm long x 30 cm wide) with a grid floor composed of rods that were two mm in diameter, spaced 15 mm apart. The front and back walls of both sets of chambers were made of Plexiglas and their side walls of aluminum. A tray containing corn cob bedding was located below the floor of each chamber. Each chamber was located in its own wooden sound and light attenuating compartment. A light source and speaker were mounted on the back wall of each compartment. The chambers were cleaned with water upon removal of a rat, and those in Context B was scented with peppermint essence (four drops onto a plastic weigh boat located in the sound attenuating cubicle). Each chamber was illuminated by an infrared light and sessions were recorded via a camera mounted on the rear wall of the cubicle and connected to a computer located in another room in the laboratory.

### Stimuli

The conditioned stimuli (CSs) were auditory and visual stimuli, counterbalanced in their roles as S1 and S2. The auditory stimulus was a 620 Hz tone of 70 dB intensity, measured at the center of the chamber (Dick Smith audiometer) and the visual stimulus was a light that flashed at 3.5 Hz with an intensity of 8 lux, measured at the center of the chamber (Lux meter). The unconditioned stimulus (US) was a 0.5s duration, 0.8 mA intensity shock delivered through the grid floor via a custom-built, constant current generator. An ammeter was used to verify the shock intensity prior to the conditioning sessions.

### Scoring and data analysis

Freezing was assessed by an experimenter blind to the experimental group. It was defined as the absence of movement except that required for breathing (Fanselow, 1980) and was measured using a time sampling procedure whereby each rat was scored every two seconds as either freezing or not freezing. The number of samples scored as freezing was divided by the total number of samples to generate a percentage freezing. Freezing data across conditioning and test sessions were analyzed using orthogonal contrasts, with group and trials being the between- and within-subjects factors, respectively. In each case, the criterion for rejection of the null hypothesis was set at alpha = 0.05.

## Methods

### Surgical procedures

Rats were surgically implanted with bilateral cannulas targeting the BLA. They were anaesthetized with 2-5% inhalant isoflurane delivered in oxygen at a flow rate of 0.5 L/min. Rats received pre-operative subcutaneous injection of the non-steroidal anti-inflammatory Carprofen (5 mg/mL, Cenvet Australia). Following the onset of stable anesthesia, rats were mounted onto a stereotaxic apparatus and 26-gauge guide cannulas were implanted into the BLA in both hemispheres (AP: +/- 2.7 mm, ML: +/- 4.9 mm, DV: 8.4 mm from Bregma). The guide cannulas were secured with four jeweler’s screws and dental cement. Immediately after surgery rats received a subcutaneous injection of prophylactic penicillin (0.05 mL, 33 mg/kg) and placed on a heating pad until recovered from the anesthetic. Rats were then returned to their home tub and allowed to recover for at least seven days prior to behavioral experiments. During recovery, rats were handled and weighed daily.

### Drug Infusion

Cycloheximide (Sigma-Aldrich) was dissolved in 70% ethanol to yield a 200 µg/µL stock solution which was diluted 1:4 with artificial cerebral spinal fluid (ACSF, Sigma Aldrich) to a final concentration of 40 µg/µL. The vehicle solution was prepared by diluting 70% ethanol 1:4 in ACSF. Infusions were carried out using injectors connected via plastic tubing to a 25 µL Hamilton syringe driven by an infusion pump. Cycloheximide or vehicle was administered at a volume of 0.5 µL bilaterally over a two min period (0.25 µL/min). The injectors were left in place for a further two min to allow for diffusion of the drug.

### Histology

Following behavioral testing rats received a lethal dose of sodium pentobarbital. Brains were removed and stored at -20°C prior to coronal sectioning (40 µm) on a cryostat. Sections containing the BLA were mounted onto glass slides and stained with cresyl violet. Cannula placements were determined with a light microscope using the BLA boundaries defined by the atlas of Paxinos and Watson (2007). Rats with cannula placements outside of these boundaries were excluded from statistical analysis. Figure 1 shows the cannula placements for all rats that were included in the study.

**Figure 1.**
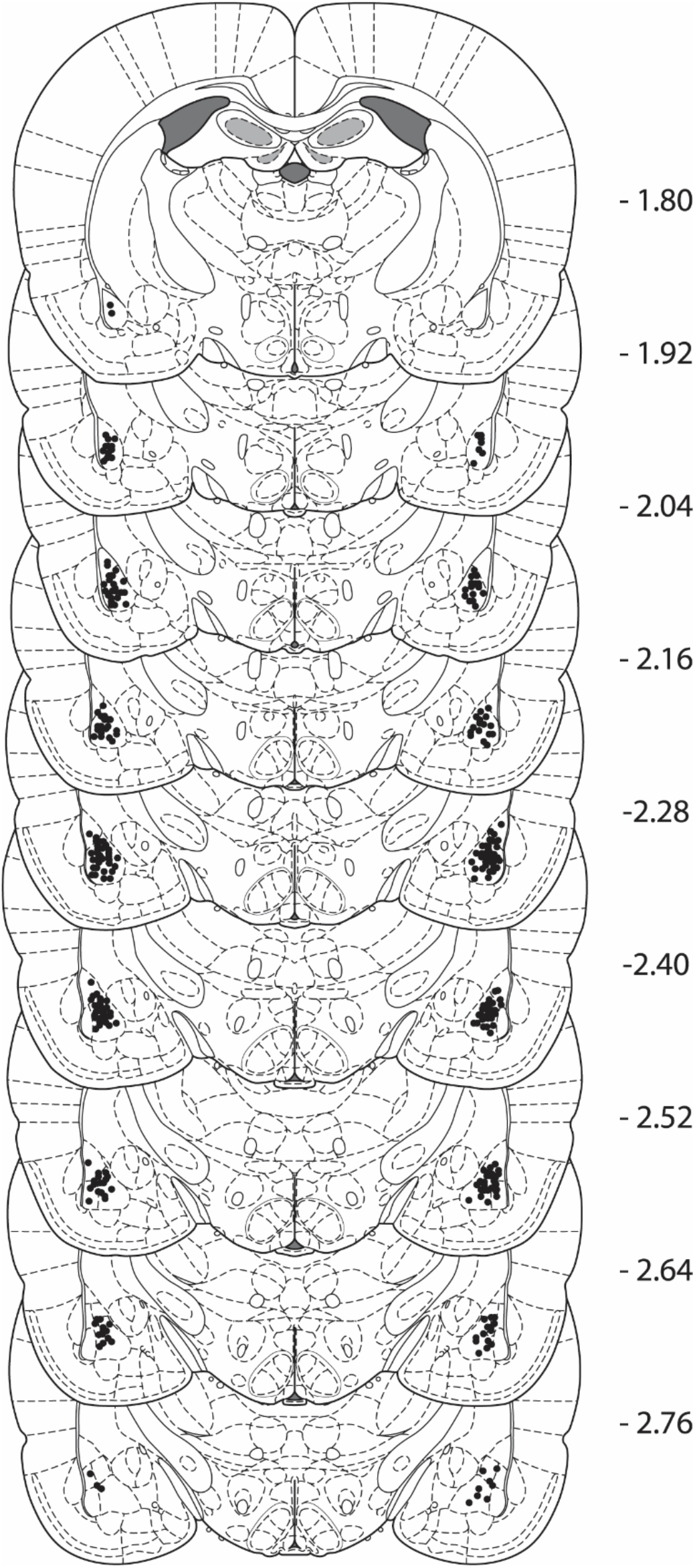
Cannula placements as verified by cresyl-violet stained sections. The black dots represent the most ventral point of the cannula track for each rat as determined using the atlas of Paxinos and Watson (2007).

## Behavioral procedures

### Experiment 1

The behavioral procedures in Experiment 1 were used in all subsequent experiments, unless otherwise indicated.

#### Context preexposure

On days 1-2, rats received twice daily exposures to the experimental chambers, one in the morning and the other approximately three hours later, in the afternoon. Each session lasted 20 min and was intended to familiarize rats with the chambers, thereby eliminating any neophobic reactions that could interfere with the detection of conditioned freezing.

#### Stage 1: S1-shock pairings (first-order conditioning)

On day 3, rats received four S1-shock pairings spaced five min apart. Each presentation of the 10 s S1 (light or tone) co-terminated with a

#### 0.5 s x 0.8 mA shock

The first S1-shock pairing occurred five min after placement in the chamber and rats were removed from the chamber one min after the last shock.

#### Context extinction

On day 4, rats received two, 20 min exposures to the context alone, once in the morning and once in the afternoon. The purpose of these sessions was to extinguish any context-elicited freezing that would otherwise mask freezing to the discrete stimuli during the subsequent stage of conditioning.

#### Stage 2: S2-S1-shock sequences (serial-order conditioning)

On day 5, rats received four S2-S1-shock sequences spaced five min apart. Offset of the 30 s S2 (tone or light) co-occurred with onset of the 10 s S1 (tone or light, respectively) which co-terminated in the 0.5 s x 0.8 mA shock. The first S2-S1-shock sequence occurred five min after placement in the chamber and rats were removed from the chamber one min after the last shock. They then received bilateral intra-BLA infusion of either vehicle or the protein synthesis inhibitor, cycloheximide.

#### Context extinction

On day 6, rats received two, 20 min exposures to the context alone, one in the morning and the other three hours later in the afternoon. They also received an additional 10 min context alone exposure on the morning of Day 7. These exposures were intended to reduce any context-elicited freezing that would obscure the test levels of freezing elicited by S2 and S1.

#### Testing

On days 7-8, rats were tested for freezing to the S2 and S1 respectively. Each test consisted of eight stimulus-alone presentations, spaced three min apart. The duration of the stimuli was identical to that used in initial conditioning: 10 s for S1 and 30 s for S2.

### Experiment 2

As in Experiment 1, rats received context familiarization on days 1 and 2, followed by S1-shock pairings on day 3. Rats in Groups P2 (paired, 2 day) and U2 (unpaired, 2 day) received the second conditioning session two days after S1-shock pairings (Day 5). Rats in Groups P14 and U14 received the second conditioning session 14 days after S1-shock pairings. Rats in the paired groups (P2 and P14) received four S2-S1-shock sequences with a five min ITI. Rats in the unpaired groups (U2 and U14) received equivalent exposures to the S2 and S1-shock but in an explicitly unpaired arrangement. Specifically, rats received four S2 alone presentation with a 2.5 min ITI, followed by four S1-shock pairings, with a 2.5 min ITI. All groups remained in the chambers for one min following the last shock. All groups received context extinction one day after S2-S1-shock sequences, followed by S2 test and S1 test as previously described in Experiment 1.

### Experiment 3

Experiment 3 was identical to Experiment 1 except that the two stages of training were separated by 2 weeks, rather than 2 days. Briefly, rats received two days of context exposure (Days 1-2), followed by S1-shock pairings (day 3). Thirteen days later (Day 16), rats received context extinction. They were then exposed to S2-S1-shock sequences (Day 17) and infused with either CHX or VEH. Rats then received context extinction (Day 18) and testing (Days 19 and 20) as previously described in Experiment 1.

### Experiment 4a

As in Experiment 3, rats received S1-shock pairings (stage 1) followed two weeks later by exposure to S2-S1-shock sequences and a BLA infusion of CHX or VEH (stage 2). The difference between the two experiments was that rats received a reminder (either S1 alone or shock alone) two days prior to the conditioning session in stage 2. Briefly, rats received two days of context exposure (Days 1-2), followed by S1-shock pairings (day 3). Twelve days later (Day 15), rats were returned to the conditioning chambers where they received two presentations of either the S1 alone (light or tone) or the shock alone with a two min ITI. The first presentation occurred five min after placement in the chamber and rats were removed two minutes after the second presentation. On day 16, rats received context extinction, followed by exposure to S2-S1-shock sequences (Day 17) and infusion of either CHX or VEH. Rats then received context extinction (Day 18) and testing (Days 19 and 20) as previously described in Experiment 1.

### Experiment 4b

The design was the same as that in Experiment 4a, except that here the reminder stimulus was presented two days after the session of S1-shock pairings (day 5) rather than two days prior to the S2-S1-shock sequences as in Experiment 4a (day 15). Across days 1-4, rats received context exposure, S1 conditioning and context extinction. On day 5, rats received two presentations of the S1 alone (light or tone counterbalanced). On day 16, rats received context extinction, followed by S2-S1-shock sequences (Day 17) and infusion of either CHX or VEH. Rats then received context extinction (Day 18) and testing (Days 19 and 20) as previously described in Experiment 1.

### Experiment 5

The next set of experiments examined the effect of shifting the context between the two stages of conditioning. In Experiment 5, rats received S1-shock training in either the same chambers used in the previous experiments (context A) or a different set of chambers (context B). Briefly, rats received two days of context exposure to both sets of chambers (Days 1-2). The order of context exposure was counterbalanced, with half of the rats being exposed to context A and then to context B and vice versa for the remainder. On day 3, rats received four S1-shock presentations in either context A (Group Same) or context B (Group Different). On day 4, rats underwent two, 20 min sessions of context extinction, one in context A and one in context B. This was intended to reduce freezing elicited by context B, and, critically, to enable detection of discrimination between the two contexts. On day 5, all rats received stage two conditioning in context A. Rats in group Same Paired and Different Paired received four S2-S1-shock sequences with a five min ITI. The remaining rats (Groups Same Unpaired and Different Unpaired) received equivalent exposures to S2 and S1-shock but in an explicitly unpaired arrangement. Specifically, rats received four S2 alone presentation with a 2.5 min ITI, followed by four S1-shock pairings, with a 2.5 min ITI. All groups remained in the chambers for one min following the last shock. On day 6, rats received context extinction in context A, followed by S2 testing (day 7) and S1 testing (day 8) in context A.

### Experiment 6

Training was similar to Group Different Paired in Experiment 5. Briefly, rats received two days of context exposure to both sets of chambers (Days 1-2). On day 3, rats received four S1-shock pairings in context B. On day 4, rats underwent two, 20 min sessions of context extinction, one in context A and one in context B. On day 5, rats received four S2-S1-shock sequences in context A, followed immediately by intra-BLA infusions of CHX or VEH. On day 6, rats received context extinction in context A, followed by S2 and S1 testing in context A on days 7 and 8 respectively.

### Experiment 7

The design was similar to Experiment 6. Rats received two days of context exposure to both context A and context B (Days 1-2), followed by S1-shock pairings in context B (Day 3). The difference between the two experiments was that, here, rats received presentations of either S1 or shock in context A prior to the session of S2-S1-shock sequences in that context. Specifically, on day 4, rats received two presentations of either S1 alone or shock alone in context A. The first stimulus presentation occurred five min after placement in the chamber and rats were removed from the chamber two minutes after the second stimulus presentation. On day 5, rats were exposed to four S2-S1-shock sequences in context A followed immediately by intra-BLA infusions of CHX or VEH. On day 6, rats received context extinction in context A, followed by testing of S2 and S1 on days 7 and 8 respectively, in context A.

## Results

### Experiment 1

The aim of Experiment 1 was to replicate previous findings (Leidl et al., 2018, Williams-Spooner et al., 2019) demonstrating that consolidation of fear to a novel S2 does not depend on *de novo* protein synthesis in the BLA when it is conditioned in compound with a previously conditioned S1. All groups successfully acquired fear to S1, with freezing increasing across its four pairings with shock (Figure 2B, *F*_(1, 23)_ = 109.44, *Fc* = 4.28, *p* < 0.001). There were no significant between-group differences (*F*_(1, 23)_ = 0.34, *Fc* = 4.28, *p* = 0.56) nor group x trend interactions (*F*_(1, 23)_ = 0.15, *Fc* = 4.28, *p* = 0.70), indicating that acquisition of fear to the S1 in stage 1 was equivalent in the two groups prior to stage 2. Similarly, acquisition of fear to the S2 was equivalent in the two groups across the S2-S1-shock sequences in stage 2 (Figure 2C). There was a significant main effect of trial (*F*_(1, 23)_ = 55.07, *Fc* = 4.28, *p* < 0.001), but no significant between-group differences (*F*_(1, 23)_ = 0.53, *Fc* = 4.28, *p* = 0.47) or group x trial interaction (*F*_(1, 23)_ = 0.06, *Fc* = 4.28, *p* = 0.81). Further there were no significant differences in the rate or level of freezing to the S1 across S2-S1-shock training (main effect of trial, *F*_(1, 23)_ = 2.66, *Fc* = 4.28, *p* = 0.12, main effect of group, *F*_(1, 23)_ = 0.07, *Fc* = 4.28, *p* = 0.79, trial x group interaction *F*_(1, 23)_ = 0.05, *Fc* = 4.28, *p* = 0.83).

**Figure 2.**
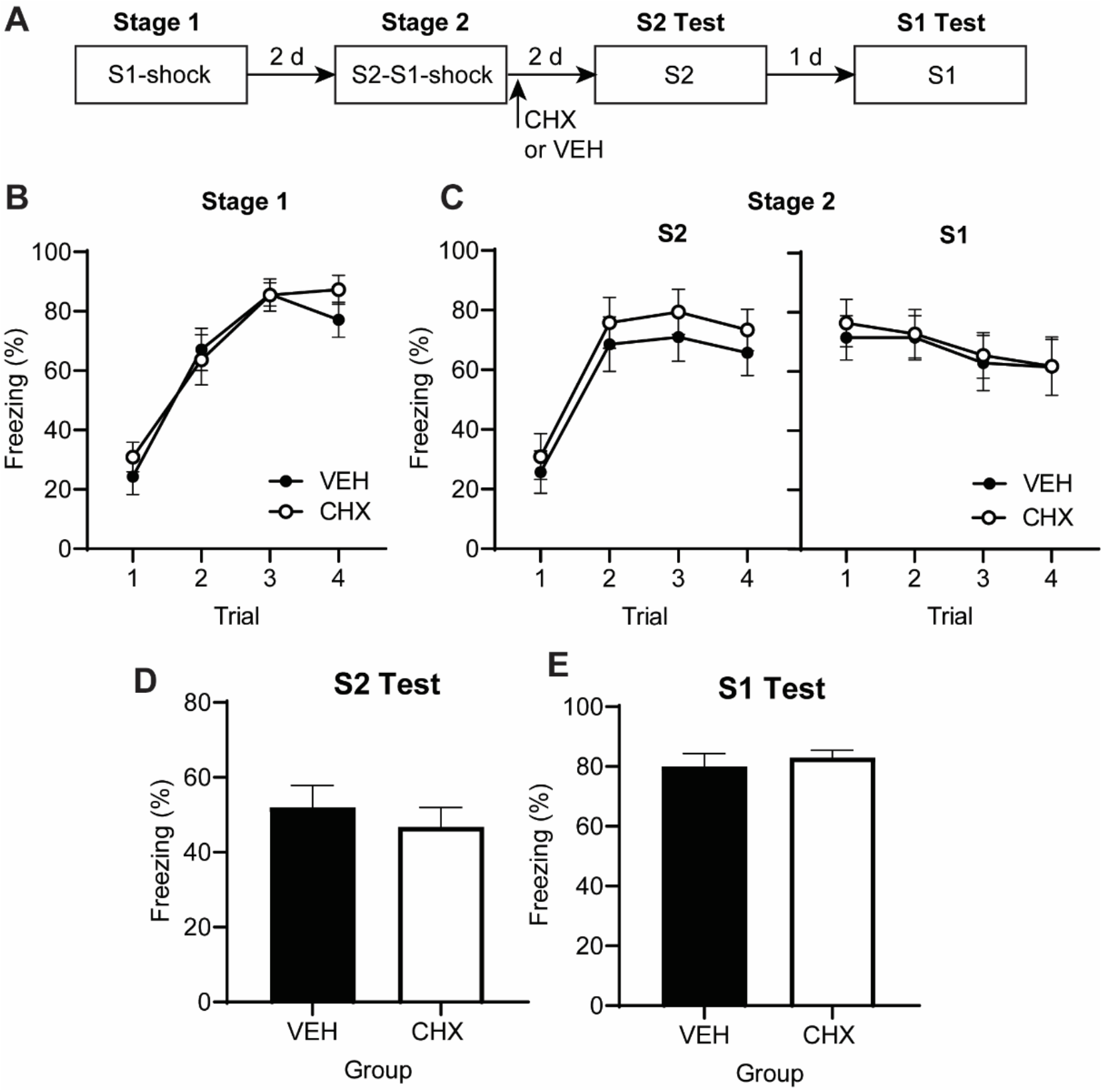
*De novo* protein synthesis in the BLA is not required to consolidate fear to the S2 when it is conditioned in compound with the already-conditioned S1. (A) Experimental timeline of conditioning and testing for rats that received intra-BLA infusions of CHX (*n* = 14) or VEH (*n* = 11). (B) Mean (±SEM) levels of freezing to presentations of the S1 during S1-shock pairings. (C) Mean (±SEM) levels of freezing to presentations of the S2 and S1 during S2-S1-shock sequences. (D-E) Mean (+SEM) freezing to test presentations of the S2 and S1 alone.

Consistent with previous findings (Leidl et al., 2018, Williams-Spooner et al., 2019, Williams-Spooner et al, 2022), both groups froze at similar levels when tested with the S2 or the S1 (Figure 2D-E). Statistical analyses confirmed there was no effect of drug on freezing levels to either the S2 (*F*_(1, 23)_ = 0.42, *Fc* = 4.28, *p* = 0.53) or the S1 (*F*_(1, 23)_ = 0.29, *Fc* = 4.28, *p* = 0.60), indicating that the intra-BLA CHX infusion did not disrupt consolidation of fear to either stimulus.

### Experiment 2

The next set of experiments examined whether the interval of time between the two conditioning experiences influences the protein synthesis requirement for consolidating the S2 fear memory. Research indicates that fear conditioning can result in a temporary increase in neural excitability (Zhou et al., 2009; Rashid et al., 2016; Josselyn and Frankland, 2018), which may alter the way neurons store subsequent information. If it is the case that consolidation of the S2 is dependent on these changes, then we would expect that inserting a delay would reverse these effects, as activity-dependent changes would have returned to baseline. Accordingly, we extended the interval between the stages of S1-shock and S2-S1-shock conditioning from the two days used previously to two weeks. We began by examining whether acquisition of fear to the S2 at a remote timepoint (14 days) was dependent on the formation of an association between S2 and/or the S1-shock. To this end, we trained four groups of rats with S1-shock pairings in stage 1. Two groups received the second stage of conditioning two days later (2 day groups) while the other two groups commenced stage two 14 days after initial conditioning (14 day groups). During this stage, one group in each pair was exposed to four S2-S1-shock sequences (Paired, Groups P2 and P14), while the other group was exposed to further S1-shock pairings and temporally separated S2 alone presentations (Unpaired, Groups U2 and U14). Rats were then tested for fear to S2 and S1. The question of interest was whether fear to the S2 was associatively mediated at both the recent and remote time points.

There was a significant increase in freezing across S1-shock pairings (Figure 3B, *F*_(1, 28)_ = 771.89, *Fc* = 4.20, *p* < 0.001). There were no significant between-group differences or group x trend interactions, indicating that acquisition of fear to the S1 was equivalent between the 2 day and 14 day groups and did not differ between paired and unpaired groups (*Fs* < 2.0, *F*_*c*_ = 4.20, *p* > 0.05).

**Figure 3.**
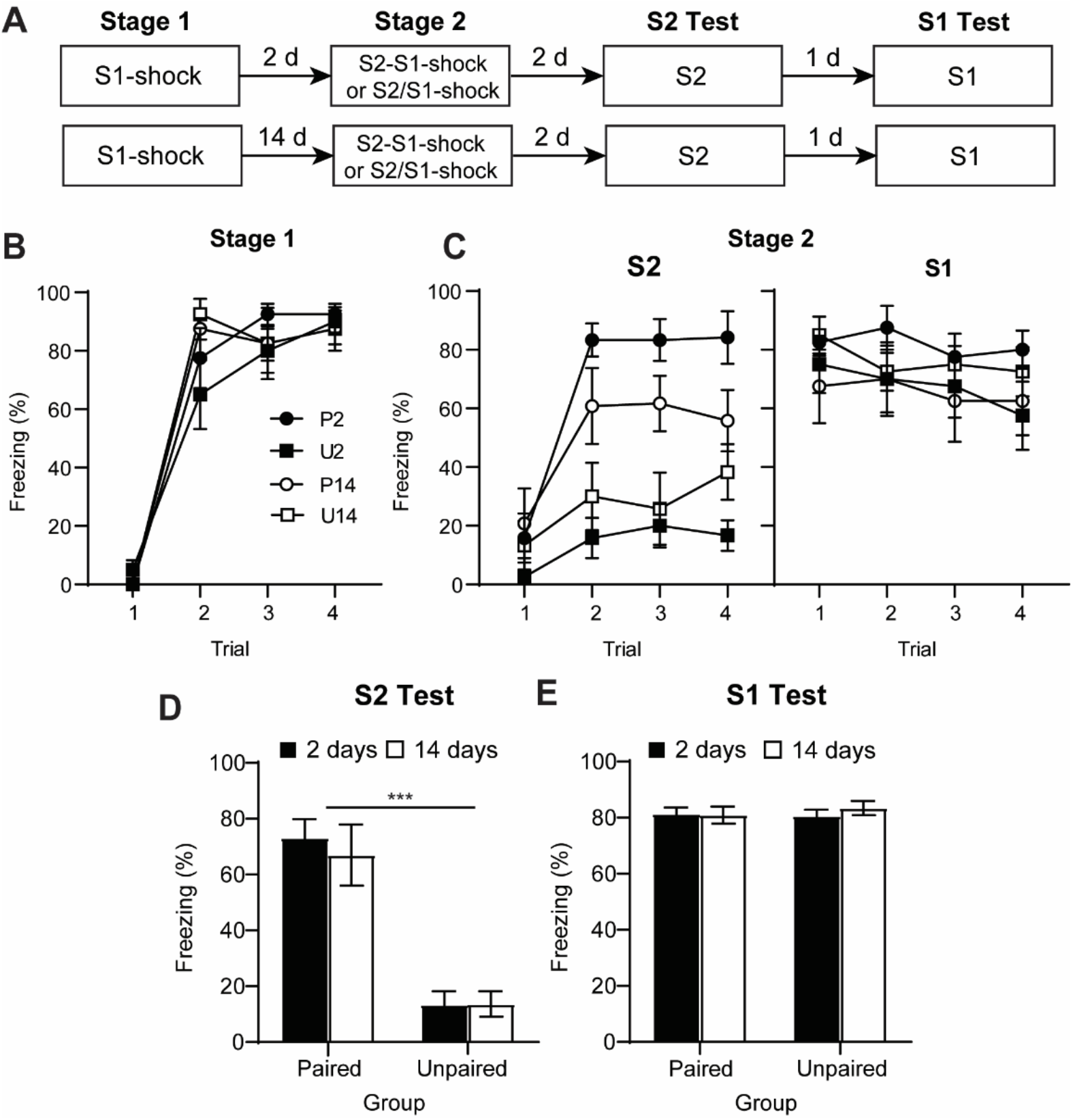
Acquisition of fear to the S2 is associatively mediated when it is conditioned two or 14 days after the prior conditioning of S1. (A) Experimental timeline of conditioning and testing. Rats received either paired or unpaired presentations of S2 and S1-shock either 2 days or 14 days after S1-shock pairings. (B) Mean (±SEM) levels of freezing to presentations of the S1 during S1-shock pairings. (C) Mean (±SEM) levels of freezing to presentations of the S2 and S1 during S2-S1-shock sequences. (D-E) Mean (+SEM) freezing to test presentations of the S2 and S1 alone. *n* = 8 for all groups.

During stage 2 (Figure 3C), rats in the paired groups displayed significantly higher freezing to the S2 than those in the unpaired groups (*F*_(1, 28)_ = 26.41, *F*_*c*_ = 4.20, *p* < 0.001). There was no overall difference in freezing levels between the 2 day and 14 day groups (*F*_(1, 28)_ = 0.07, *F*_*c*_ = 4.20, *p* = 0.80) and no significant time interval x pairing interaction (*F*_(1, 28)_ = 4.13, *F*_*c*_ = 4.20, *p* = 0.052). Averaging across groups, there was a significant increase in freezing across conditioning trials (*F*_(1, 28)_ = 52.37, *F*_*c*_ = 4.20, *p* < 0.001). This effect was driven by the paired groups, which increased freezing across S2-S1-shock sequences, while freezing in the unpaired groups remained low, resulting in a significant paired x trend interaction (*F*_(1, 28)_ = 10.67, *F*_*c*_ = 4.20, *p* < 0.01). There was no significant time interval x trend interaction (*F*_(1, 28)_ = 1.61, *F*_*c*_ = 4.20, *p* = 0.215), although there was a significant time x paired x trend interaction (*F*_(1, 28)_ = 4.34, *F*_*c*_ = 4.20, *p* < 0.05), indicating that the increase in freezing to the S2 across trials was more pronounced for rats in Group P2 relative to those in group P14. Freezing to the S1 across stage two decreased across trials (*F*_(1, 28)_ = 5.60, *F*_*c*_ = 4.20, *p* < 0.05), likely reflecting an increase in fear-elicited escape behaviors. Freezing levels to the S1 did not differ between the groups and there were no significant interactions involving any between- or within-subjects factors (*Fs* < 2.2, *F*_*c*_ = 4.20).

At test (Figure 3D-E), freezing levels to the S2 were higher in the paired than the unpaired groups (*F*_(1, 28)_ = 105.22, *F*_*c*_ = 4.20, *p* < 0.001) . In contrast, there was no significant difference in levels of freezing to the S1 (*F*_(1, 28)_ = 0.026, *F*_*c*_ = 4.20, *p* = 0.87). In both cases, there was no significant effect of the time interval between conditioning stages, or time x context interaction (*Fs* < 2.4, *F*_*c*_ = 4.20). Together, this indicates that conditioning to the S2 at both the recent and remote time point is conditional upon its pairing with S1-shock, rather than generalization of fear from the conditioned S1.

### Experiment 3

This experiment examined whether increasing the interval between the two conditioning experiences to two weeks alters the requirement for protein synthesis in consolidating the S2 fear memory. Rats received S2-S1-shock conditioning 14 days after initial S1-shock conditioning. Immediately after S2-S1-shock conditioning, rats were infused with either CHX or VEH. They were then tested for fear to both the S2 and the S1.

Rats acquired fear to the S1 in stage 1 and the S2 in stage 2 (Figure 4B-C). In stage 1, freezing to the S1 increased across its pairing with shock (*F*_(1, 22)_ = 82.64, *Fc* = 4.30, *p* < 0.001). There was an unexpected between-group difference (*F*_(1, 22)_ = 12.45, *Fc* = 4.30, *p* < 0.005), with the rats that would be assigned to the CHX group freezing more than those assigned to the VEH group. There was no significant trial x drug interaction (*F*_(1, 22)_ = 0.47, *Fc* = 4.30, *p* = 0.50) indicating that the difference between groups was consistent across S1-shock pairings. Importantly, this difference was not maintained across the S2-S1-shock sequences in stage 2, with both groups showing similar levels of freezing to the S1 (*F*_(1, 22)_ = 0.03, *Fc* = 4.30, *p* = 0.86). Statistical analysis of freezing to the S2 across the sequences confirmed a main effect of trial (*F*_(1, 22)_ = 42.2, *Fc* = 4.3, *p* < 0.001) but no main effect of group (*F*_(1, 22)_ = 0.04, *Fc* = 4.30, *p* = 0.85) or group x trial interaction (*F*_(1, 22)_ = 1.47, *Fc* = 4.30, *p* = 0.24).

**Figure 4.**
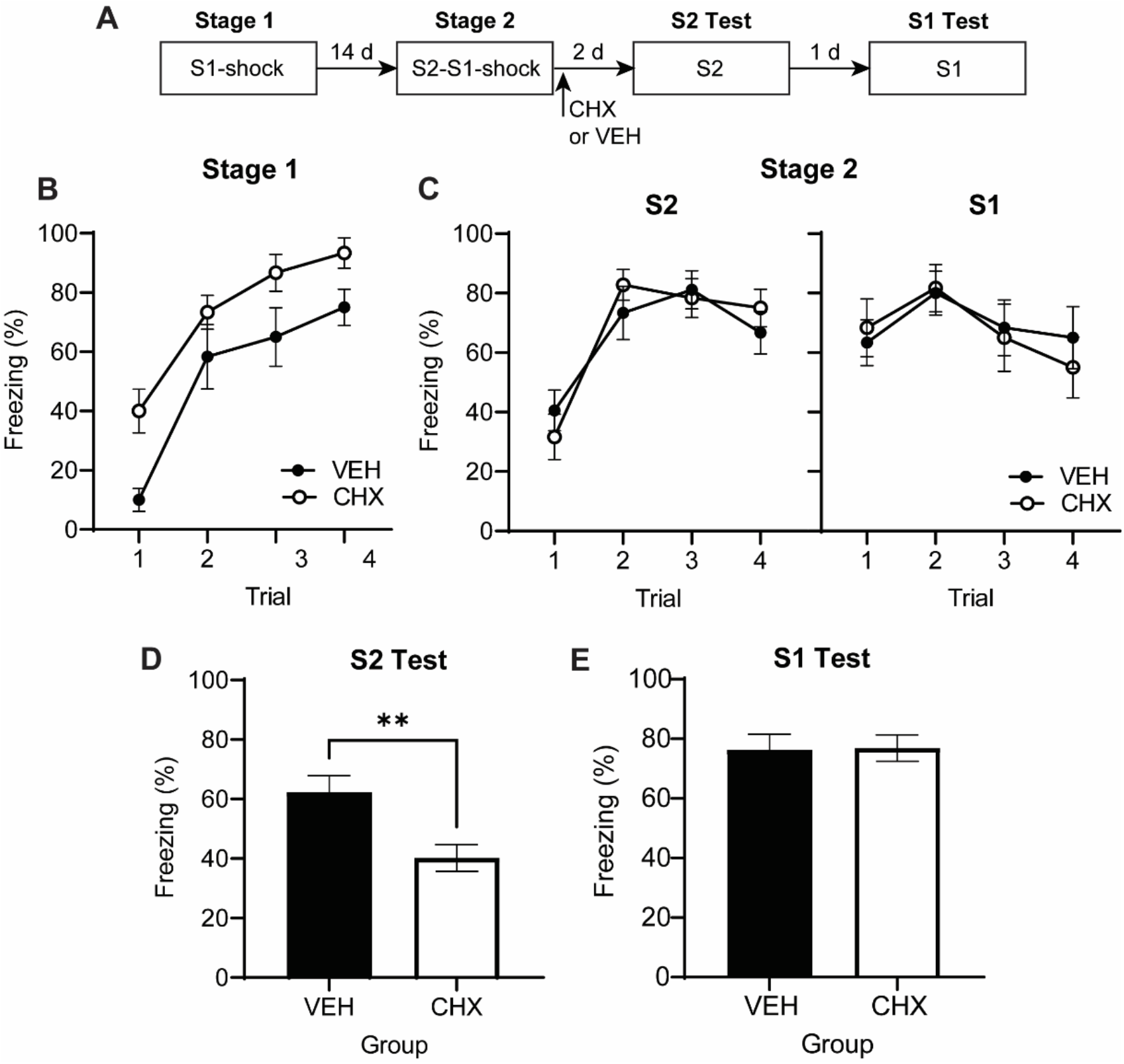
A two-week interval reinstates the protein synthesis requirement for consolidating fear to the S2 in the BLA. (A) Experimental timeline of conditioning and testing for rats that received intra-BLA infusions of CHX (*n* = 12) or VEH (*n* = 12). (B) Mean (±SEM) levels of freezing to presentations of the S1 across its pairings with shock. (C) Mean (±SEM) levels of freezing to presentations of the S2 and S1 across the S2-S1-shock sequences. (D-E) Mean (+SEM) freezing across test presentations of S2 alone and S1 alone.

During S2 test (Figure 4D), rats in the VEH group froze significantly more than rats in the CHX group, indicating that conditioning of the S2 had been disrupted by the BLA infusion of CHX (*F*_(1, 22)_ = 9.76, *Fc* = 4.30, *p* < 0.01). In contrast, there was no significant difference between groups in freezing to the S1 (Figure 4E, *F*_(1, 22)_ = 0.008, *Fc* = 4.30, *p* = 0.93). In combination with Experiment 1, these results indicate that the time between stages influences the requirement for *de novo* protein synthesis in consolidating the S2 fear memory.

### Experiment 4

Consolidation of the S2 fear memory required protein synthesis in the BLA when S2 was conditioned (S2-S1-shock sequences in stage 2) two weeks after the prior conditioning of S1 (S1-shock pairings in stage 1). Experiments 4a and 4b examined whether reminding rats of their prior experience in stage 2 would render consolidation of S2 insensitive to the effects of protein synthesis inhibition. We hypothesized that a reminder would functionally bridge the gap between the two experiences, and hence, consolidation of the S2 fear memory would no longer require *de novo* protein synthesis in the BLA. In order to test this possibility, rats received S1-shock pairings (stage 1) and two weeks later, S2-S1-shock sequences, followed by an intra-BLA infusions of CHX or VEH (stage 2). The experiments differed with respect to when the reminder occurred. In Experiment 4a, rats received the reminder (either two presentations of the S1 alone or two shocks) two days prior to S2-S1-shock sequences in stage 2. In experiment 4b, they received the reminder 12 days prior to S2-S1-shock sequences in stage 2 (i.e., two days after the S1-shock pairings).

### Experiment 4a

The two vehicle infused groups exhibited equivalent freezing across S1-shock pairings in stage 1, S2-S1-shock sequences in stage 2, and during testing with S2 and S1. Therefore, they were combined into a single vehicle control group and the data are presented for rats in three groups: those for which stage 2 was preceded by the S1 reminder and followed by the BLA cycloheximide infusion (Group CHX S1); those for which stage 2 was preceded by the shock reminder and followed by the BLA cycloheximide infusion (Group CHX shock); and those in the composite control group for which stage 2 was preceded by one of the reminder treatments (S1 or shock) and followed by a BLA infusion of vehicle (VEH).

All groups acquired fear to the S1 in stage 1 and to the S2 in stage 2 (Figure 5A-B). Specifically, freezing to S1 increased across its pairings with shock in stage 1 (*F*_(1, 27)_ = 134.8, *Fc* = 4.2, *p* < 0.001). There were no significant differences between infusion type (*F*_(1, 27)_ = 0.32, *Fc* = 4.2, *p* = 0.58) or reminder type (*F*_(1, 27)_ = 1.63, *Fc* = 4.2, *p* = 0.21), and no significant group x trend interactions (*Fs* < 0.55, *Fc* = 4.17, *p* > 0.05). Similarly, freezing to S2 increased across the four S2-S1-shock sequences in stage 2 (*F*_(1, 27)_ = 132.4, *Fc* = 4.2, *p* < 0.001); and, again, there were no significant differences between infusion type (*F*_(1, 27)_ = 0.004, *Fc* = 4.2, *p* = 0.95) or reminder type (*F*_(1, 27)_ = 0.41, *Fc* = 4.2, *p* = 0.53), and no significant group x trend interactions (*Fs* < 2.6, *Fc* = 4.17, *p* > 0.05). Freezing to the S1 remained consistently high across these sequences (main effect trend *F*_(1, 27)_ =1.25, *Fc* = 4.2, *p* = 0.23) and there was no significant effect of infusion condition (*F*_(1, 27)_ = 0.23, *Fc* = 4.2, *p* = 0.59) or reminder condition (*F*_(1, 27)_ = 0.04, *Fc* = 4.2, *p* = 0.85), and no significant group x trend interactions (*Fs* < 1.5, *Fc* = 4.17, *p* > 0.05).

**Figure 5.**
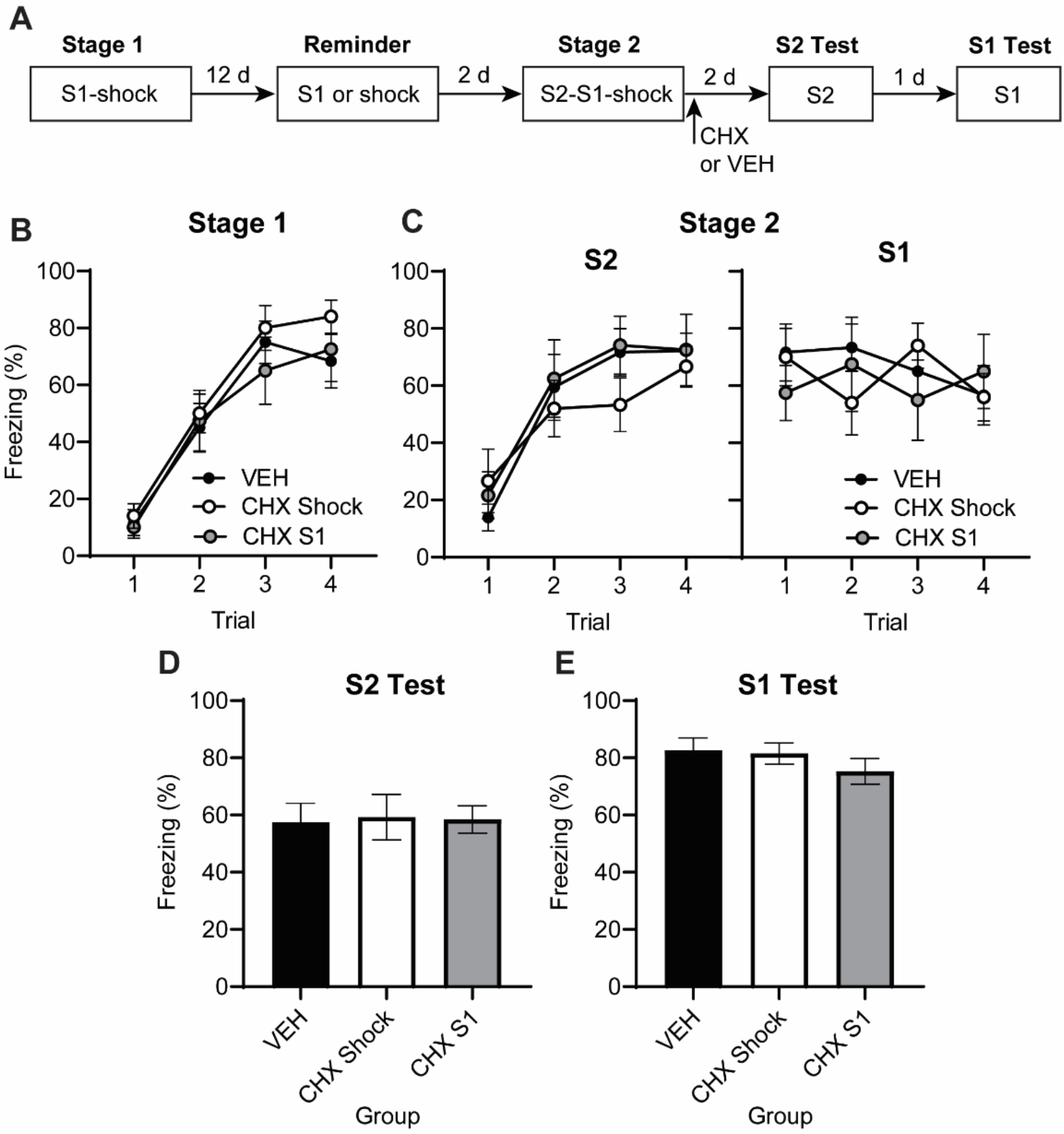
Reminding rats of their prior experience reverses the effect of a time delay on the protein synthesis requirement for consolidating fear to the S2. (A) Experimental timeline of conditioning, reminder and testing for rats that received intra-BLA infusions of CHX or VEH. (B) Mean (±SEM) levels of freezing to S1 across its pairing with shock; to S2 and S1 across S2-S1-shock sequences (C); and to S2 (D) and S1 (E) at test. VEH, *n* = 13, CHX shock, *n* = 10, CHX S1 *n* = 8.

Freezing levels at test are presented in Figure 5D-E. Inspection of the figure suggests that levels of freezing were equivalent across groups, regardless of infusion condition or reminder type. This was confirmed via statistical analyses which revealed no significant effect of infusion condition (S1, *F*_(1, 27)_ = 0.74, *Fc* = 4.2, *p* = 0.53; S2, *F*_(1, 27)_ = 0.03, *Fc* = 4.2, *p* = 0.40) or reminder type (S1, *F*_(1, 27)_ = 0.92, *Fc* = 4.2, *p* = 0.35; S2, *F*_(1, 27)_ = 0.006, *Fc* = 4.2, *p* = 0.94) on freezing levels to either the S1 or the S2. Thus, reminding rats of their stage 1 experience (S1-shock) two days prior to conditioning in stage 2 (S2-S1-shock) renders consolidation of the S2 fear memory independent of protein synthesis in the BLA.

### Experiment 4b

Experiment 4b was identical to Experiment 4a, except that the reminder was presented 12 days (instead of two days) prior to serial-order conditioning in stage 2. As we previously found that S1 and shock were equally effective as reminders, in the current experiment all rats received only S1 reminders.

All groups successfully acquired freezing responses to the S1 in stage 1 and to the S2 in stage 2 (Figure 6B-C). Specifically, there was an increase in freezing to the S1 across its pairings with shock in stage 1 (*F*_(1, 23)_ = 196, *Fc* = 4.3, *p* < 0.001). This increase occurred at a similar rate among rats in groups CHX and VEH (*F*_(1, 23)_ = 0.16, *Fc* = 4.3, *p* = 0.69) and, overall, there were no significant between-group differences (*F*_(1, 23)_ = 0.001, *Fc* = 4.3, *p* = 0.98). Similarly, there was an increase in freezing to S2 across the S2-S1-shock sequences in stage 2 (*F*_(1, 23)_ = 72.9, *Fc* = 4.3, *p* < 0.001). The overall level of freezing to S2 did not differ between the CHX and VEH groups (S2, *F*_(1, 23)_ = 0.31, *Fc* = 4.3, *p* = 0.58) and there was no significant group x trial interaction (*F*_(1, 23)_ = 1.1, *Fc* = 4.3, *p* = 0.30). Freezing to S1 across the S2-S1-shock sequences did not differ between groups (*F*_(1, 23)_ = 0.14, *Fc* = 4.3, *p* = 0.71) or change across trials (*F*_(1, 23)_ = 0.26, *Fc* = 4.3, *p* = 0.62), and there was no significant group x trial interaction (*F*_(1, 23)_ = 0.14, *Fc* = 4.3, *p* = 0.71).

**Figure 6.**
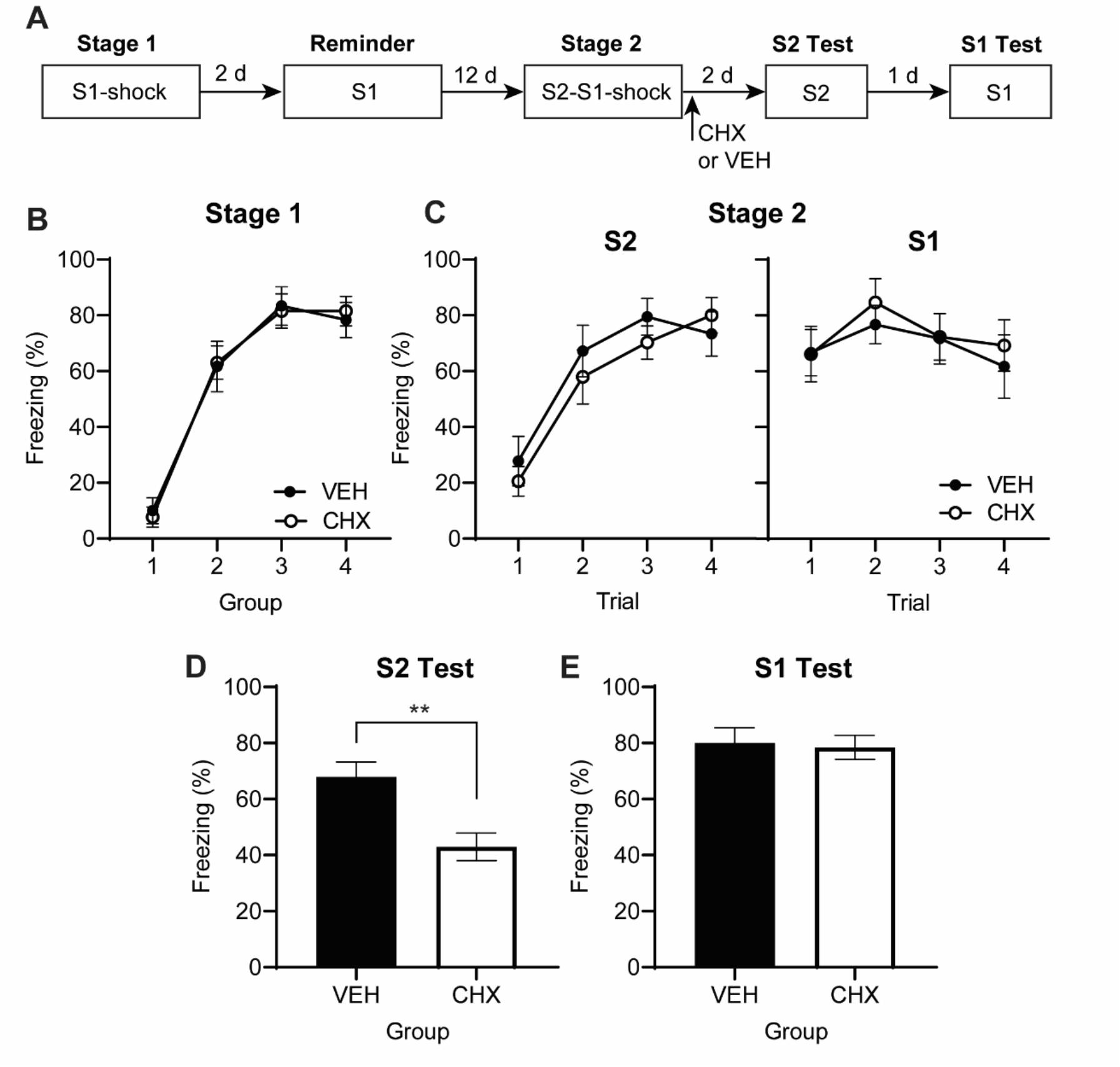
Presenting the reminder 12 days prior to stage 2 spares the effect of a time delay on the protein synthesis requirement for consolidating fear to S2. (A) Experimental timeline of conditioning, reminder and testing for rats that received intra-BLA infusions of CHX (*n* = 13) or VEH (*n* = 12). (B) Mean (±SEM) levels of freezing to presentations of the S1 during S1-shock pairings. (C-D) Mean (±SEM) levels of freezing to presentations of the S2 and S1 during S2-S1-shock pairings. (E-F) Mean (+SEM) levels of freezing to presentations of the S1 and S2 during testing.

At test (Figure 6D-E), rats in group CHX froze significantly less to the S2 than those in group VEH (*F*_(1, 23)_ = 11.9, *Fc* = 4.3, *p* < 0.01) while both groups showed similar levels of freezing to the S1 (*F*_(1, 23)_ = 0.05, *Fc* = 4.3, *p* = 0.83). Together with the results of the previous experiment, this shows that the effect of a reminder on how fear to S2 is consolidated depends on when the reminder is presented. When the reminder is temporally distant from the session of S2-S1-shock sequences, consolidation of fear to S2 requires protein synthesis in the BLA. By contrast, when the reminder is presented close to the session of S2-S1-shock sequences, consolidation of fear to S2 ceases to require protein synthesis in the BLA.

### Experiment 5

Experiments 2-4 examined the effect of a shift in temporal context on the requirement for BLA protein synthesis in consolidating the S2 fear memory. Experiments 5-7 examine the effects of a shift in physical context between the S1-shock pairings in stage 1 and the S2-S1-shock sequences in stage 2. We began by examining whether acquisition of fear to S2 in the different context was dependent on formation of an association between S2 and/or the S1-shock. To this end, we trained four groups of rats with S1-shock pairings in stage 1. Two groups then received stage 2 in the same context (Groups Same) while the remaining two groups received stage 2 in a second different context (Groups Different). For one group in each of these pairs, stage 2 consisted in four S2-S1-shock sequences (Groups Same Paired and Different Paired). For the remaining group in each pair, stage 2 training consisted in S1-shock pairings and temporally separated S2 alone presentations (Groups Same Unpaired and Different Unpaired). Rats were then tested for fear to S2 and S1 in the stage 2 context. The question of interest was whether acquisition of fear to the S2 was associatively mediated in the same and different context conditions.

All groups successfully acquired fear to S1 in stage 1, with freezing increasing across the four conditioning trials (Figure 7B, *F*_(1, 31)_ = 61.722, *p* < 0.001). There were no significant between-group differences or group x trend interactions, indicating that acquisition of fear to S1 was equivalent between groups prior to manipulations of context and did not differ between paired and unpaired groups (*F* < 1.5, *F*_*c*_ = 4.160).

**Figure 7.**
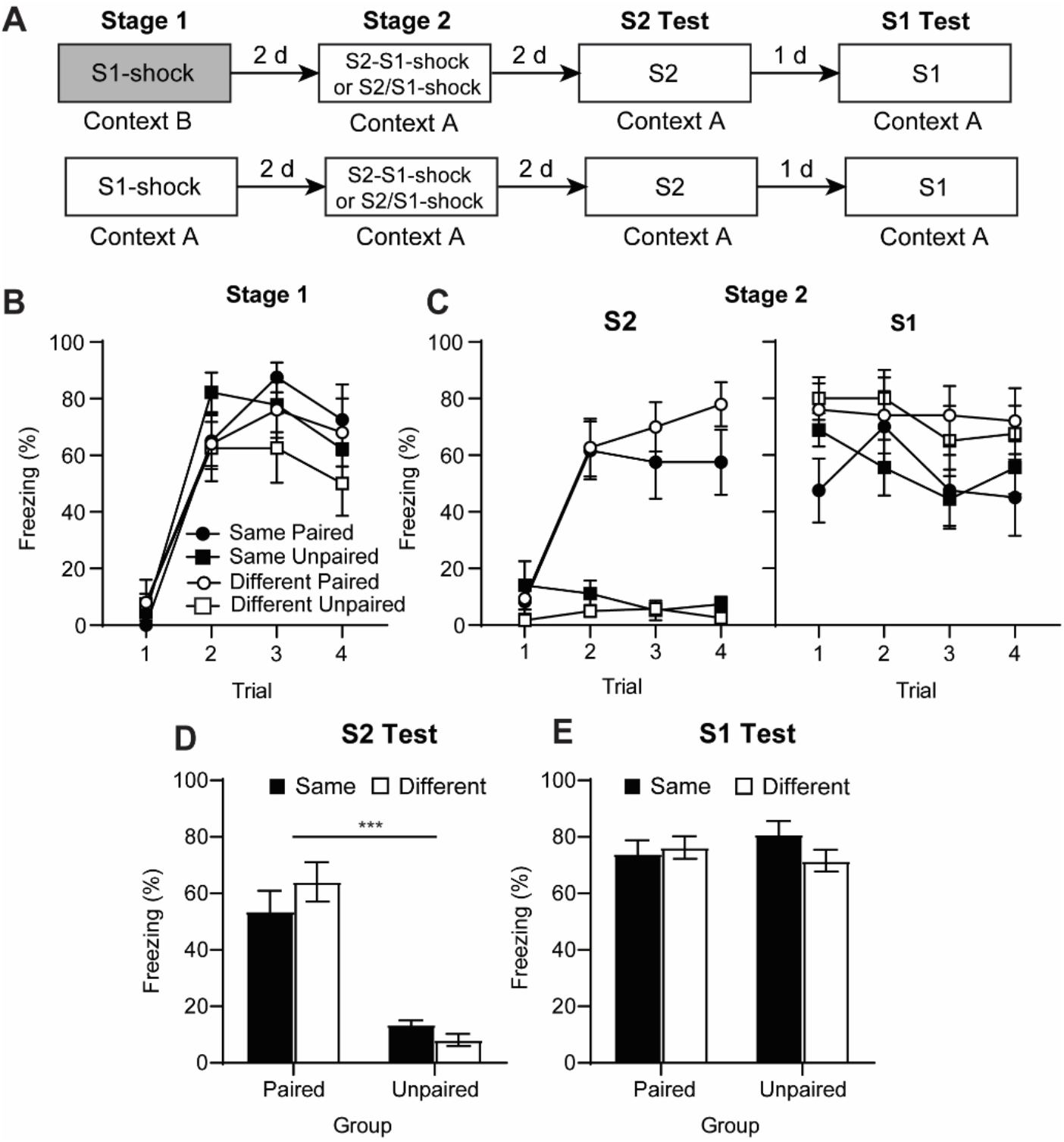
Fear to the S2 is associatively mediated, even when it is conditioned in a different context to that of the prior S1-shock pairings. In stage 1, rats received S1-shock pairings in either context B or context A. In stage 2, rats received either paired or unpaired presentations of S2 and S1-shock. (B) Mean (±SEM) levels of freezing to presentations of the S1 during S1-shock pairings. (C) Mean (±SEM) levels of freezing to presentations of the S2 and S1 during S2-S1-shock sequences. (D-E) Mean (+SEM) freezing to test presentations of the S2 and S1 alone. Group Same Paired, *n* = 8, Same Unpaired, *n* = 9, Different Paired, *n* = 10, Different Unpaired, *n* = 8.

In stage 2, freezing developed to S2 among rats that received S2-S1-shock sequences but there was relatively little freezing to S2 in rats that received the S2 explicitly unpaired from S1-shock pairings (Figure 7C, *F*(1, 31) = 60.555, *F*_*c*_ = 4.160, *p* < 0.001). Averaging across groups, freezing increased across conditioning trials (*F*_(1, 31)_ = 34.888, *F*_*c*_ = 4.160, *p* < 0.001). This effect was driven by performance in the paired groups, which increased across presentations of S2, whilst freezing in the unpaired groups remained low, leading to a significant trend x pairing interaction (*F*_(1, 31)_ = 48.497, *F*_*c*_ = 4.160, *p* < 0.001). There was no significant effect of context or context x pairing interaction (*Fs* < 3.2, *F*_*c*_ = 4.160), indicating that freezing to S2 was similar regardless of whether the context was the same or different to that in stage 1. This was not due to a failure to discriminate between the two contexts, as the context-alone exposures revealed significantly higher freezing in the context where the S1-shock pairings had occurred than in the other context (Figure 1E, *F*_(1, 68)_ = 17.910, *F*_*c*_ = 3.982, *p* < 0.001). Unexpectedly, freezing to the S1 across stage 2 was significantly higher in the different groups compared to the same groups (*F*_(1, 31)_ = 5.368, *F*_*c*_ = 4.160, *p* < 0.05). This freezing did, however, remain stable across the stage (*F*_(1, 31)_ = 3.412, *F*_*c*_ = 4.160, *p* = 0.074) and there were no significant differences between paired and unpaired groups or trend x pairing interactions (*F*_s_ < 3.5, *F*_*c*_ = 4.160).

At test (Figure 7D-E), freezing levels to S2 were higher in the paired than the unpaired groups (*F*_(1, 31)_ = 79.90, *F*_*c*_ = 4.160, *p* < 0.001). In contrast, there were no significant differences in freezing to S1 (*Fs* < 2, *F*_*c*_ = 4.161). In both cases, there was no significant effect of context or context x pairing interaction (*Fs* < 2.5, *F*_*c*_ = 4.161) indicating that shifting the context between stages 1 and 2 did not affect overall performance to S1 or S2. Together, these results indicate that acquisition of fear to the S2 was conditional on its close temporal contiguity with S1-shock pairings; and unaffected by whether the S2-S1-shock sequences were administered in the same or different context to that of the prior S1-shock pairings.

### Experiment 6

Experiment 6 examined the effect of a context shift between stages 1 and 2 on the protein synthesis requirement for consolidating fear to the S2. To this end, rats received S1-shock pairings in context B (stage 1) and then received S2-S1-shock sequences in context A (stage 2). Immediately following the session containing the S2-S1-shock sequences, rats received a BLA infusion of CHX or VEH. Rats were then tested for fear to the S2 and S1 in context A.

Both groups successfully acquired freezing to S1 in stage 1 (Figure 8B). There was a significant increase in freezing to the S1 across its pairing with shock (*F*_(1, 32)_ = 92.40, *Fc* = 4.15, *p* < 0.001) with no between-group differences in overall levels of freezing or group x trial interaction (main effect group, *F*_(1, 32)_ = 0.001, *Fc* = 4.15, *p* = 0.98, group x trend interaction, *F*_(1, 32)_ = 0.018, *Fc* = 4.15, *p* = 0.89). During the subsequent context exposure session, rats showed significantly higher freezing in the context where S1-shock pairings had occurred (context B) than in the other context (context A) where rats would undergo all remaining conditioning and testing (*F*_(1, 32)_ = 34.47, *Fc* = 4.15, *p* < 0.001). However, examination of the freezing levels in the two contexts revealed individual differences in context discrimination, with a small number (*n* = 5) of rats in the CHX group showing higher levels of freezing in the non-shocked context. We assumed that the effect of the context shift manipulation would only be effective in those rats that successfully discriminated between the two contexts. Therefore, the test data for the cycloheximide infused rats was divided into those that could discriminate (‘discriminators’) and those that failed to do so (‘generalizers’). Discriminators were defined as those rats that showed higher levels of freezing in the shock context (context B) than the alternative context (context A), whereas the generalizers froze more in the non-shocked context (context A).

**Figure 8.**
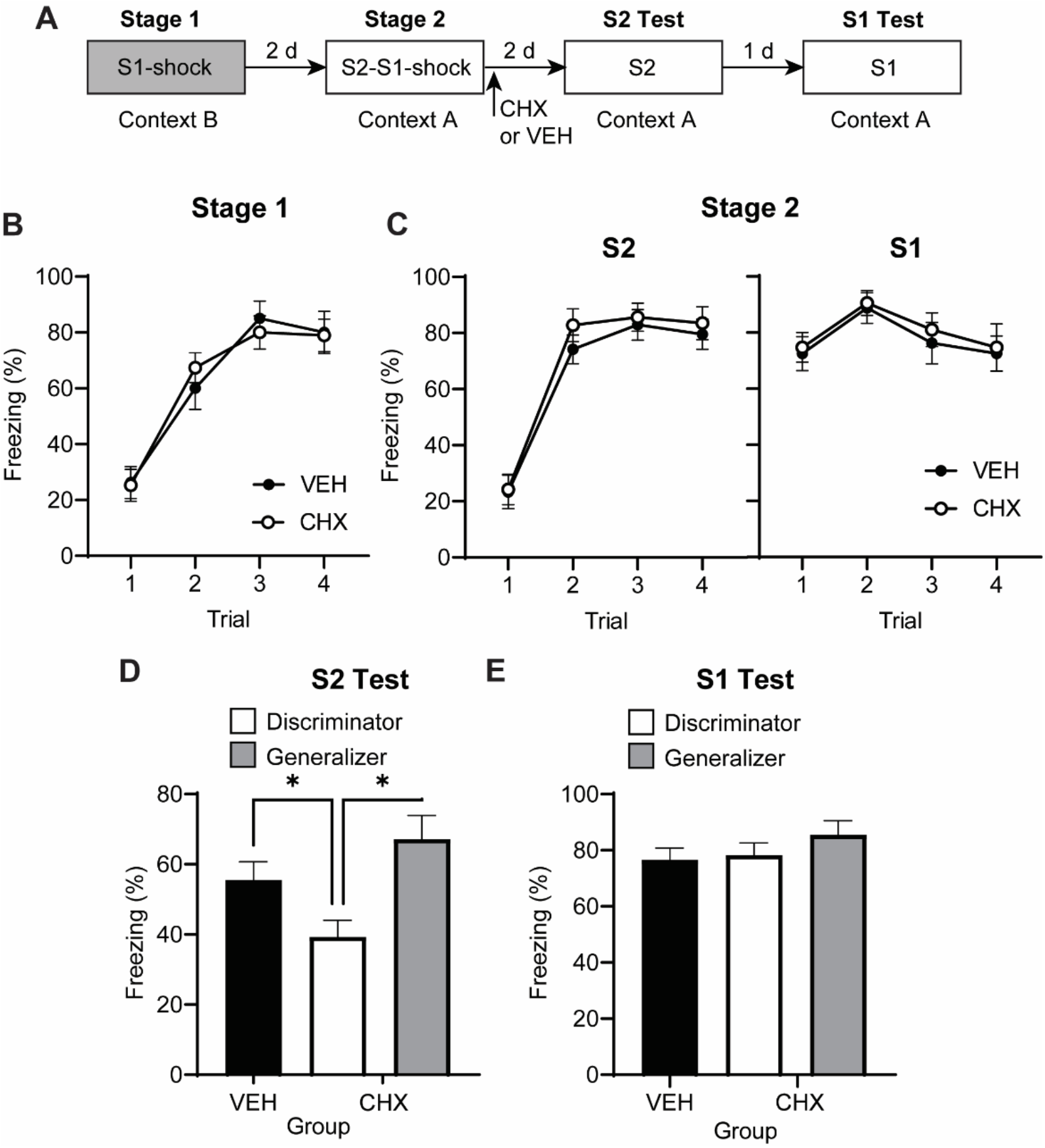
When the two stages of training occur in different physical contexts *and rats can tell these contexts apart*, the protein synthesis requirement for consolidating fear to S2 is reinstated. (A) Experimental timeline of conditioning, reminder and testing for rats that received an intra-BLA infusion of CHX or VEH. (B) Mean (±SEM) levels of freezing to presentations of the S1 during S1-shock pairings. (C) Mean (±SEM) levels of freezing to presentations of the S2 and S1 during S2-S1-shock sequences. (D-E) Mean (+SEM) freezing to test presentations of the S2 and S1 alone. VEH, *n* = 15, CHX discriminators, *n* = 14, CHX generalizers *n* = 5.

Across the S2-S1-shock sequences in context A, rats increased freezing to the S2 (*F*_(1, 32)_ = 160.2, *Fc* = 4.15, *p* < 0.001), while their freezing to the S1 remained stable (*F*_(1, 32)_ = 0.53, *Fc* = 4.15, *p* = 0.47). There were no significant differences between VEH and CHX groups in overall freezing to either the S2 (Figure 8C, *F*_(1, 32)_ = 0.27, *Fc* = 4.15, *p* = 0.61) or the S1 (*F*_(1, 32)_ = 0.25, *Fc* = 4.15, *p* = 0.62) and no significant trial x group interactions (S2, *F*_(1, 32)_ = 0.005, *Fc* = 4.15, *p* = 0.94; S1, *F*_(1, 32)_ = 0.09, *Fc* = 4.15, *p* = 0.77).

The test levels of freezing to the S2 and S1 are shown in Figure 8, panels D and E. Inspection of the figure suggests that rats in the CHX discriminator group froze less to the S2 than rats in the CHX generalizer group and VEH controls. This impression was confirmed by the statistical analyses, which showed that the overall levels of freezing to the S2 was significantly lower in the CHX discriminator group than the CHX generalizer group and VEH controls groups (*F*_(1, 31)_ = 10.19, *Fc* = 4.15, *p* < 0.01). Further, there was no significant difference in freezing between the CHX generalizers and the VEH infused controls (*F*_(1, 31)_ = 1.48, *Fc* = 4.15, *p* = 0.23) indicating that the effect of CHX was contingent upon the rat’s ability to discriminate between the two contexts. All groups froze at similar levels to the S1, with no significant between group differences (CHX discriminator vs other groups, *F*_(1, 31)_ = 0.23, *Fc* = 4.15, *p* = 0.64, CHX generalizer vs. VEH, *F*_(1, 31)_ = 1.19, *Fc* = 4.15, *p* = 0.28). Taken together, these results show that a BLA infusion of cycloheximide disrupts conditioning to the S2 when there is a shift in the physical context between S1-shock pairings in stage 1 and S2-S2-shock sequences in stage 2. However, this disruption is dependent upon the rats being able to successfully identify that they are in a different context.

### Experiment 7

This experiment examined whether reminding rats of their stage 1 experience in the stage 2 context would alter the protein synthesis requirement (in the BLA) for consolidating fear to S2. Rats received S1-shock pairings in context B. Forty-eight hours later they received a shock or a non-reinforced presentation of S1 (tone or light) in the different context, A; and 48 hours after that, they were exposed to S2-S1-shock sequences in context A. The session containing the sequences was followed by an intra-BLA infusion of either CHX or VEH. Finally, rats were tested for freezing to S2 and S1 in context A. If reminding rats of their stage 1 experience in the stage 2 context (prior to the S2-S1-shock sequences) renders the A and B contexts functionally equivalent, consolidation of fear to S2 should be unaffected by the BLA cycloheximide infusion. That is, the reminder should reverse or block the context shift effect observed in the previous experiment.

Five rats were excluded from the statistical analyses due to a failure to discriminate between the two contexts after S1-shock pairings in context B. Based on the results of Experiment 6, a failure to discriminate between the contexts would render consolidation of the S2 fear memory insensitive to the effects of cycloheximide, which is the expected consequence of presenting the conditioned S1 in context A before serial-order conditioning. Therefore, these rats were excluded from the statistical analyses.

The two vehicle infused groups exhibited equivalent levels of freezing across all stages of training and testing and were therefore combined into a single vehicle control group. The data is presented with respect to three groups: those for which stage 2 was preceded by an S1 reminder and followed by a BLA infusion of CHX (Group CHX S1); those for which stage 2 was preceded by a shock reminder and followed by a BLA infusion of CHX (Group CHX shock); and the composite control group that received either a S1 or shock reminder followed by a BLA infusion of vehicle (VEH).

Both CHX and VEH groups acquired freezing to S1 across its pairings with shock in context B and to S2 across the S2-S1-shock sequences in context A (Figure 9B-C). Specifically, there was an increase in freezing to S1 across its pairings with shock in stage 1 (*F*_(1, 30)_ = 206.31, *Fc* = 4.17, *p* < 0.001) and to S2 across S2-S1-shock sequences in stage 2 (*F*_(1, 30)_ = 135.60, *Fc* = 4.17, *p* < 0.001). There was no significant difference in freezing between the infusion conditions (S1, *F*_(1, 30)_ = 0.2, *Fc* = 4.171, *p* = 0.658; S2, *F*_(1, 30)_ = 0.944, *Fc* = 4.17, *p* = 0.51), reminder type (S1, *F*_(1, 30)_ = 1.12, *Fc* = 4.171, *p* = 0.30; S2, *F*_(1, 30)_ = 1.07, *Fc* = 4.17, *p* = 0.49), or group x trend interactions (*Fs* < 0.6, *Fc* = 4.17, *p* > 0.05) indicating that freezing to the S1 and the S2 did not differ prior to drug manipulations. Freezing to the S1 across the S2-S1-shock sequences did not differ between infusion conditions (*F*_(1, 30)_ = 1.592, *Fc* = 4.17, *p* = 0.22) or reminder type (*F*_(1, 30)_ = 1.54, *Fc* = 4.17, *p* = 0.70), and did not change across trials (*F*_(1, 30)_ = 0.257, *Fc* = 4.17, *p* = 0.616). There were no significant group x trial interactions (*Fs* < 0.2, *Fc* = 4.17, *p* > 0.05).

**Figure 9.**
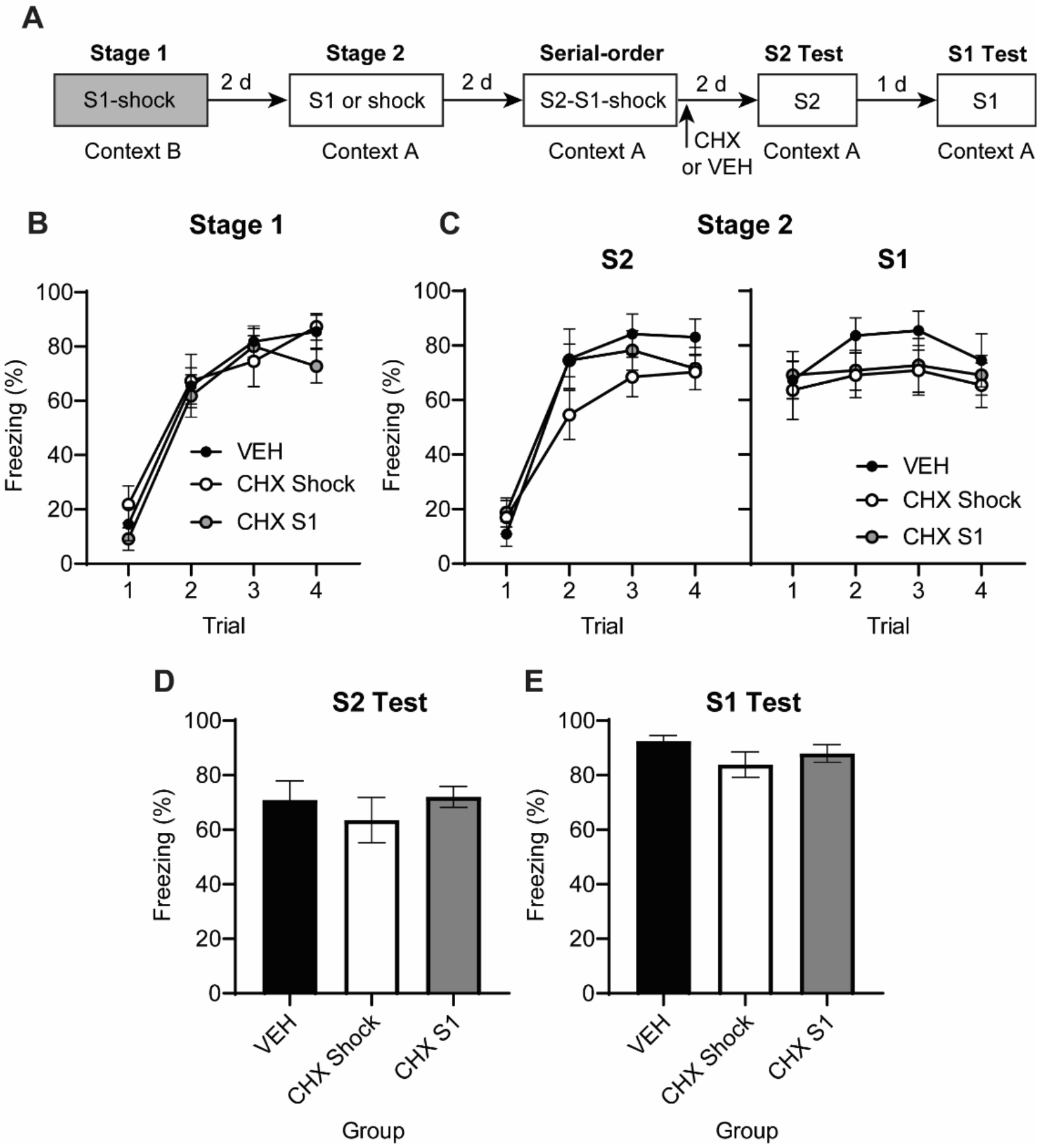
Re-exposing rats to S1 or shock in the stage 2 conditioning context alleviates the protein synthesis requirement for consolidating fear to S2. (A) Experimental timeline of conditioning, reminder and testing for rats that received intra-BLA infusions of CHX or VEH. (B) Mean (±SEM) levels of freezing to presentations of the S1 during S1-shock pairings. (C) Mean (±SEM) levels of freezing to presentations of the S2 and S1 during S2-S1-shock sequences. (D-E) Mean (+SEM) freezing to test presentations of the S2 and S1 alone. VEH, *n* = 11, CHX shock, *n* = 11, CHX S1 *n* = 11.

At test (Figure 9D-E), there was no significant difference between CHX and VEH groups in freezing to either S2 (*F*_(1, 30)_ = 0.146, *Fc* = 4.17, *p* = 0.71) or S1 (*F*_(1, 30)_ = 2.38, *Fc* = 4.17, *p* = 0.13). There was also no significant effect of reminder type on levels of freezing to the S2 (*F*_(1, 30)_ = 0.81, *Fc* = 4.17, *p* = 0.37) or S1 (*F*_(1, 30)_ = 0.69, *Fc* = 4.17, *p* = 0.41). Together, this indicates that presentation of the conditioned S1 or shock in the context (A) where S2 would be conditioned alleviated the protein synthesis requirement for consolidating fear to S2.

## Discussion

Protein synthesis in the BLA is not required to consolidate the memory of a new dangerous experience when it is similar to a past experience (Leidl et al., 2018; Williams-Spooner et al., 2019). The present study examined whether the timing and location of the new dangerous experience (relative to the past experience) affects the protein synthesis requirement for its consolidation in the BLA. In each experiment, rats were trained using a two-stage fear conditioning protocol in which they received S1-shock pairings in stage one and S2-S1-shock sequences in stage two. The latter stage was immediately followed by a BLA infusion of the protein synthesis inhibitor, cycloheximide or vehicle. Finally, all rats were tested with S2 and S1.

The first set of experiments showed that the protein synthesis requirement for consolidating fear to S2 varies with the interval between the two training stages. Specifically, consolidation of fear to S2 did not require protein synthesis in the BLA when the two training stages occurred 48 hours apart (Experiment 1), as previously reported (Leidl et al., 2018; Williams-Spooner et al., 2019); but did require this synthesis when the interval between the two stages was increased to 14 days (Experiment 3). However, under the latter circumstances, the protein synthesis requirement was additionally influenced by reminders of the stage one training prior to stage two: the BLA cycloheximide infusion disrupted fear to S2 among rats that were re-exposed to S1 twelve days prior to stage two (Experiment 4b); but had no effect among rats that were re-exposed to the S1 (or shock) just 48 hours prior to stage two (Experiment 4a). Thus, within the BLA, the protein synthesis requirement for consolidating fear to S2 is *reinstated* with time since the prior conditioning of S1; but this reinstatement can be blocked or reversed by a reminder of the prior S1 conditioning shortly before the conditioning to S2. When this occurs, consolidation of fear to S2 remains (or, is again) independent of protein synthesis in the BLA.

The second set of experiments showed that the protein synthesis requirement for consolidating fear to S2 also changes when the second training stage is conducted in a different physical context. Here, rats were exposed to the initial S1-shock pairings (stage one) in one context and, 48 hours later, to the S2-S1-shock sequences (stage two) in a second, distinct context. The results showed that fear to the S2 was disrupted by a BLA infusion of cycloheximide among rats that *could* successfully discriminate between the contexts (Experiment 6). However, this effect of cycloheximide was not observed when rats could *not* discriminate between the contexts (Experiment 6); and when, 48 hours prior to stage two, rats were reminded of their prior S1-shock experience in the different context (Experiment 7). Thus, the impact of a context shift on the protein synthesis requirement for consolidating fear to S2 appears to depend on the history or ‘meaning’ of the context in which S2 is conditioned. When the contexts differ with respect to their association with the dangerous cues, the context shift results in protein synthesis-dependent consolidation of fear to S2; but when both contexts are associated with dangerous cues, consolidation of fear to S2 does not require protein synthesis in the BLA.

The finding that a delay and context shift influence consolidation of fear to S2 in a similar way suggests that their effects are mediated by a common mechanism. One way of thinking about this is that: 1) time is a gradually changing temporal context (Bouton, 2002); and 2) context plays a hierarchical or occasion-setting role in signaling whether a new situation is the same or different to a prior experience (Bouton and Swartzentruber, 1986; Fraser and Holland, 2019). However, it is important to note that the form of occasion-setting implied by the present findings differs from that which has been previously described, whereby the context (or another CS) regulates the retrieval of specific CS-US relations. That is, the novel implication of the present findings is that context does not only influence the processes by which memories are retrieved and expressed in behavior: it also influences the substrates by which new information is consolidated in the brain, or at least in the BLA.

How does the brain compare situations and detect similarity/dissimilarity and what are the consequences of this comparison? A large literature implicates the hippocampus in context processing and computing the overlap between past and present experience based on convergent cortical inputs (Marr, 1971; Kim and Fanselow, 1992; Phillips and LeDoux, 1992; Eichenbaum, 2015). The hippocampus has reciprocal connections with the amygdala and has been shown to regulate context-dependent learning and synaptic changes in the amygdala (Pitkänen et al., 2000; Orsini et al., 2011; Ferrara et al., 2019). Accordingly, the hippocampus may use features of the environment to identify the context as the same or different and transfer this information to the BLA. When both experiences occur in the same physical and temporal context, the hippocampus identifies the current situation as the same as that experienced previously and, consequently, fear that develops to S2 is linked to the established fear of S1. However, when the two experiences occur in different physical/temporal contexts, the mismatch is identified by the hippocampus and fear to S2 is stored independently of fear to S1.

One question that remains to be addressed is how fear to S2 is consolidated independently of protein synthesis in the BLA. We propose that, when the novel S2 is presented in sequence with the already-conditioned S1 and shock, fear to S2 is acquired through activation of the same BLA neurons that encoded/retrieved the prior S1-shock memory: hence, it does not require the full suite of molecular changes for its consolidation. However, with a change in physical or temporal context, the second experience is allocated to a separate population of BLA neurons and protein synthesis is again required for consolidation of fear to the S2. This proposal is supported by research demonstrating that dangerous events that occur close in time, or within the same physical context, are more likely to share overlapping neural signatures compared to events that are separated in space and time (Guzowski et al., 1999; Cai et al., 2016; Rashid et al., 2016). Applied to the present findings, initial conditioning of the S1 may lead to a temporary increase in neuronal excitability and plasticity related products in a population of BLA neurons (Josselyn and Frankland, 2018). When rats are exposed to S2-S1-shock sequences 48 hours later, fear to S2 is consolidated via molecular changes in BLA neurons that encoded the prior S1-shock memory. When rats are exposed to S2-S1-shock sequences 14 days later or in a different context, fear to S2 is consolidated via *de novo* changes in a non-specific population of BLA neurons. However, re-exposure to the S1 prior to the session containing the S2-S1-shock sequence re-excites the BLA neurons that encoded the prior S1-shock memory: hence, fear to S2 is again consolidated via molecular changes in these neurons (for evidence that a retrieval cue can return engram cells to an excited state, see Pignatelli et al. (2019)).

Finally, is important to recognize parallels between the present findings and those reported in previous studies examining the involvement of NMDA receptors in the encoding of a second dangerous experience (Crestani et al., 2018; Williams-Spooner et al., 2022). These studies show that encoding of a second dangerous experience does not require NMDA receptors in the BLA or hippocampus when it occurs within 2-3 days of a similar dangerous experience, but does require NMDA receptor activity when the two experiences are separated by 14-40 days or occur in different physical contexts. Together with the present results, these findings imply that a prior dangerous experience does not just influence the encoding *or* consolidation of a new dangerous experience: it influences both. That is, the way that a new dangerous experience is processed in the BLA and hippocampus is *fundamentally* different when it is similar to a past experience.

In summary, the present study has shown that the protein synthesis requirement for consolidating a new dangerous experience in the BLA is regulated by its similarity to a past experience. Specifically: 1) protein synthesis in the BLA is not required for consolidating fear to S2 when S2-S1-shock sequences occur in the same physical and temporal context as a prior session of S1-shock pairings; 2) the protein synthesis requirement for consolidating fear to S2 is reinstated when the two training stages are separated by a delay or occur in different physical contexts; and 3) the effects of the delay and context shift can be blocked by reminding animals of the prior S1-shock pairings. Thus, the substrates of learning and memory in the mammalian brain are not fixed and immutable. Instead, the way we learn about and remember the details of current events reflects the degree to which they match the details of our past experience, which includes information about context and time; and is influenced by recent reminders of our past experience.

## Acknowledgements

This work was supported by Australian Research Council Discovery Grants to NMH and RFW (DP200102969) and RFW (DP220103650).

